# The chlamydial transcriptional regulator Euo is a key switch in cell form developmental progression but is not involved in the committed step to the formation of the infectious form

**DOI:** 10.1101/2024.05.17.594781

**Authors:** Cody R. Appa, Nicole A. Grieshaber, Hong Yang, Anders Omsland, Sean McCormick, Travis J. Chiarelli, Scott S. Grieshaber

## Abstract

Bacteria in the genus *Chlamydia* are a significant health burden world wide. They infect a wide range of vertebrate animals including humans and domesticated animals. In humans, *C. psittaci* can cause zoonotic pneumonia while *C. pneumoniae* causes a variety of respiratory infections. Infections with *C. trachomatis* cause ocular or genital infections. All chlamydial species are obligate intracellular parasites of eukaryotic cells and are dependent on a complex infection cycle that depends on transitions between specific cell forms. This cycle consists of cell forms specialized for host cell invasion, the Elementary Body (EB), and a form specialized for intracellular replication, the Reticulate Body (RB). In addition to the EB and RB there is a transitionary cell form that mediates the transformation between the RB and the EB, the Intermediate Body (IB). In this study we ectopically expressed the regulatory protein Euo and showed that high levels of expression resulted in reversible arrest of the development cycle. The arrested chlamydial cells were trapped phenotypically at an early IB stage of the cycle. These cells had exited the cell cycle but had not shifted gene expression from RB-like to IB/EB-like. This arrested state was dependent on continued expression of Euo. When ectopic expression was reversed, Euo levels dropped in the arrested cells which led to the repression of native Euo expression and the resumption of the developmental cycle. Our data are consistent with a model where Euo expression levels impact IB maturation to the infectious EB but not the production of the IB form.

**Importance:** Bacterial species in the Chlamydiales order infect a variety of vertebrate animals and are a global health concern. They cause various diseases in humans, including genitital and respiratory infections. The bacteria are obligate intracellular parasites that rely on a complex infectious cycle involving multiple cell forms. All species share the same life cycle, transitioning through different states to form the infectious elementary body (EB) to spread infections to new hosts. The Euo gene, encoding a DNA binding protein, is involved in regulating this cycle. This study showed that ectopic expression of Euo halted the cycle at an early stage. This arrest depended on continued Euo expression. When Euo expression was reversed, the developmental cycle resumed. Additionally, this study suggests that high levels of Euo expression affects the formation of the infectious EB, but not the production of the cell form committed to EB formation.

## Introduction

The bacteria in the Chlamydiales order are intracellular parasites of eukaryotic cells (1). Within the Chlamydiales order, the genus *Chlamydia* contains the causative agents of a number of important pathogens of humans. *C. psittaci* causes zoonotic infections resulting in pneumonia, while *C. pneumoniae* is a human pathogen responsible for respiratory disease. Biovars of *C. trachomatis (Ctr)* are the causative agents of trachoma, the leading cause of preventable blindness worldwide, as well as sexually transmitted infections with the potential to cause pelvic inflammatory disease and infertility. Irrespective of the resulting disease, all chlamydial species share the same obligate intracellular life cycle and developmental cell forms, and completion of this developmental cycle is central to chlamydial pathogenesis. The developmental cycle includes the replicating cell form called the reticulate body (RB), the infectious non replicating form called the elementary body (EB), and an intermediate form (IB) that mediates the transition from the RB to the EB (2). This complex cycle does not produce typical growth culture dynamics of lag, log and stationary phases but instead asynchronously progresses through a number of distinct cell type transitions that ultimately result in amplification and dissemination of the pathogen.

*Ctr* must transition through the RB and IB phenotypic states to form the infectious EB. To regain infectivity, chlamydial cells must exit the cell cycle, halt DNA replication, express type III effectors required for the next infection, decrease in size and reorganize the chromosome into the characteristic condensed EB nucleoid. The factors that regulate the changes in gene expression underlying these dramatic phenotypic shifts are poorly defined. The *euo* gene (Early Upstream ORF) encodes a helix-turn-helix DNA binding protein that is highly conserved across the Chlamydiota phylum (3). In *C. trachomatis,* Euo has been shown to repress genes expressed late in the chlamydial developmental cycle (4, 5). Additionally, Euo has been shown biochemically to bind to many sites on the chlamydial chromosome that correlate with both repression and activation of gene expression (4, 5). In this study we ectopically expressed the Euo protein and assessed its effects on completion of the developmental cycle.

Our data show that ectopic expression of Euo arrested the developmental cycle at an early IB stage. The arrested chlamydial cells had exited the cell cycle and did not continue chromosomal replication. Upon repression of ectopic Euo expression, the developmental cycle was reinitiated and infectious EBs were produced. This recovery was biphasic with an early fast recovery due to the arrested cells immediately reinitiating EB maturation, followed by a slower replication dependent recovery suggestive of a return to RB dependent production of IBs that mature into EBs. Additionally, the data also showed that Euo acts in a feed forward loop in part controlling the switch to IB/EB gene expression that ultimately creates the infectious form.

## Results

### Ectopic expression of Euo leads to a reduction in EB production

The developmentally regulated protein Euo is among the earliest genes expressed post EB to RB germination during chlamydial infection (5). It has been shown to have DNA binding and transcriptional repression capabilities, preferentially binding A/T rich regions of the genome (6). Prior studies determined that Euo acts to repress “late genes” in the chlamydial developmental cycle, “late genes” being defined as those upregulated during the process of EB maturation at the end of the intracellular developmental cycle (5). To understand the role of Euo on the chlamydial developmental cycle, we expressed Euo ectopically in *C. trachomatis* under the control of a riboswitch (L2-E-Euo-FLAG) (Fig 1A). This construct was tested for regulation by theophylline (Tph) using western blotting and a protein of the correct size was visible only in the induced sample (Fig. S1).

**Figure 1:**
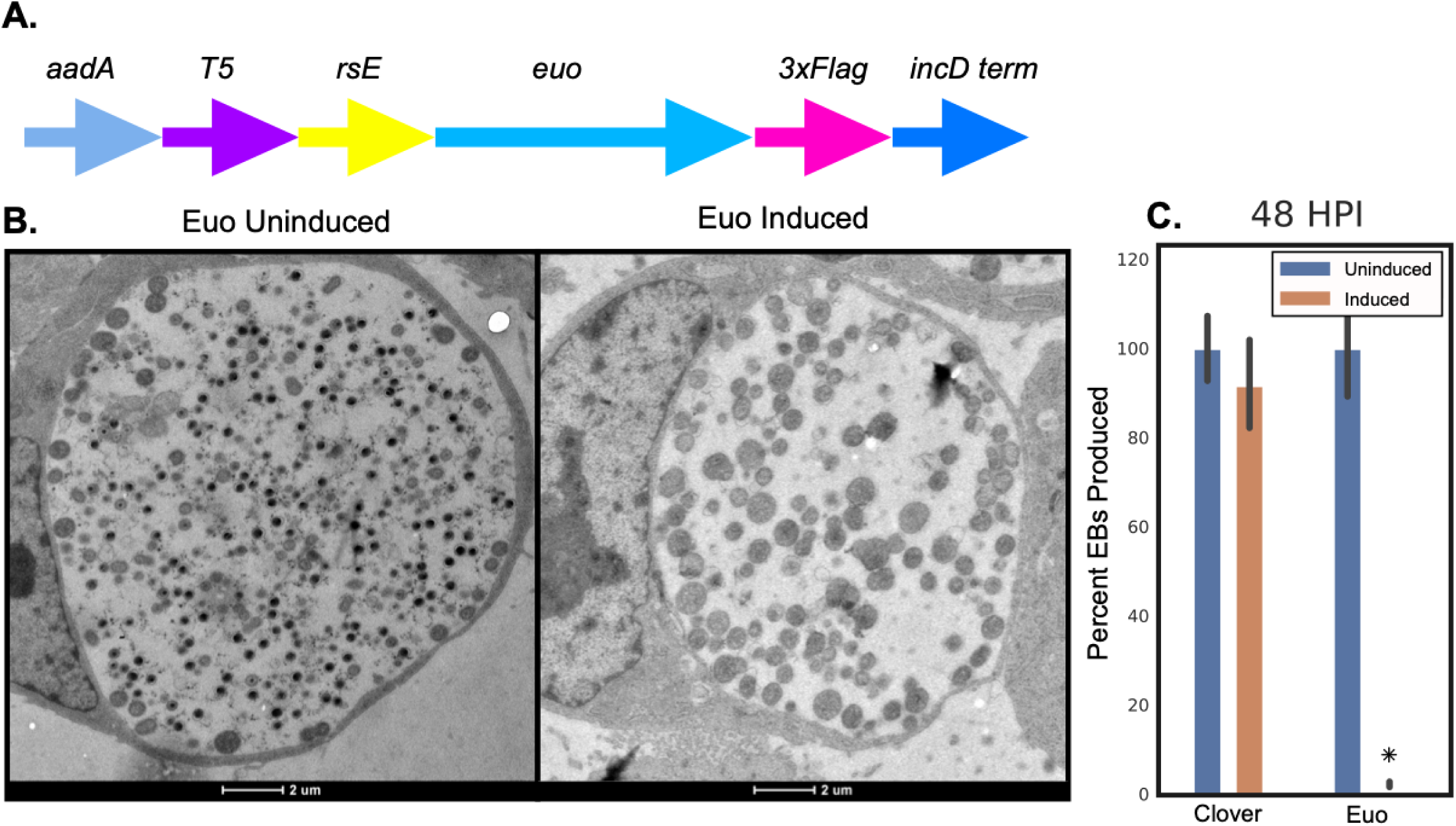
Ectopic expression of Euo-FLAG. A) Schematic of the E-Euo-FLAG construct which consisted of the streptomycin/spectinomycin resistance gene (aadA), T5-lac promoter (T5), riboE riboswitch (rsE), the chlamydial *euo* gene (*euo)*, and an in-frame 3xFLAG tag. B) Electron micrographs of the inclusions from cells infected with L2-E-Euo-FLAG and induced for Euo expression using Tph as compared to vehicle control. Euo was induced at 15 hpi and samples fixed at 30 hpi. C) Inclusion forming unit (IFU) reinfection assay from Cos-7 cells infected with either L2-E-Clover-FLAG (control) or L2-E-Euo-FLAG induced at 16 hpi with 0.5 mM Tph compared to uninduced controls. Isolation of EBs and re-infection of Cos-7 monolayers to determine the production of infectious progeny occurred at 48 hpi. Asterisks = p < 0.01.

Cos-7 cells were infected with L2-E-Euo-FLAG and Euo expression was induced with 0.5mM Tph at 15 hpi. At 30 hpi cells were fixed and imaged using transmission electron microscopy (TEM). Euo Induction resulted in dramatic phenotypic changes of *Chlamydia*. Untreated cultures contained a diverse cell population, including RBs (large cells) and EBs (small electron dense cells), within the inclusions (Fig 1B, Euo Uninduced). The chlamydial cells in the Tph-treated cultures were much more homogeneous and were missing obvious EBs (Fig. 1B, Euo Induced). Reinfection assays also demonstrated a dramatic and statistically significant decrease in infectious progeny produced in the Tph-treated cell population (Fig. 1C).

### Ectopic Euo expression arrested the developmental cycle

To visualize cell form-specific promoter activity and assess the effects of Euo ectopic expression on the kinetics of the chlamydial developmental cycle, we cloned the dual promoter reporter cassette *hctB*prom-mkate2_*euo*prom-clover (BmEc) (7) into the E-Euo-FLAG plasmid resulting in E-Euo-BmEc (Fig 2A). This plasmid was transformed into *Ctr*, creating the strain L2-E-Euo-BmEc. As published previously, *euo*prom+ cells are RBs while *hctB*prom+ cells are EBs (7). Cos-7 cells were infected with L2-E-Euo-BmEc and induced for Euo expression at 18 hpi and cultures fixed 12 hours later at 30 hpi (Fig. 2B). Induction of Euo expression resulted in an increase in green *euo*prom+ cells and a dramatic decrease in red *hctB*prom+ cells when compared to the uninduced samples, suggesting that Euo expressing cells were arrested in the cycle at a step prior to EB formation (Fig. 2B). As the arrested cell population was positive for *euo*prom activity and negative for *hctB*prom activity, this data also suggested the cells were arrested in the RB form. We employed fluorescent in situ hybridization (FISH) to determine the effects of Euo ectopic expression on the mRNA expression of three other developmentally regulated genes, *incD*, *hctA* and *tarp*. IncD is an inclusion membrane protein that is expressed early during infection (RB gene), HctA is a DNA nucleoid associated protein expressed at an intermediate time point (IB gene), while Tarp is a type III secretion system (T3SS) effector that is expressed late in the developmental cycle (EB gene) (7–9). Staining of the Euo overexpressing cells using custom FISH probes showed that Euo ectopic expression resulted in a dramatic up-regulation of the RB gene *incD*, and conversely a dramatic down-regulation of both the IB gene, *hctA,* and the EB gene, *tarp* (Fig. 2C).

**Figure 2:**
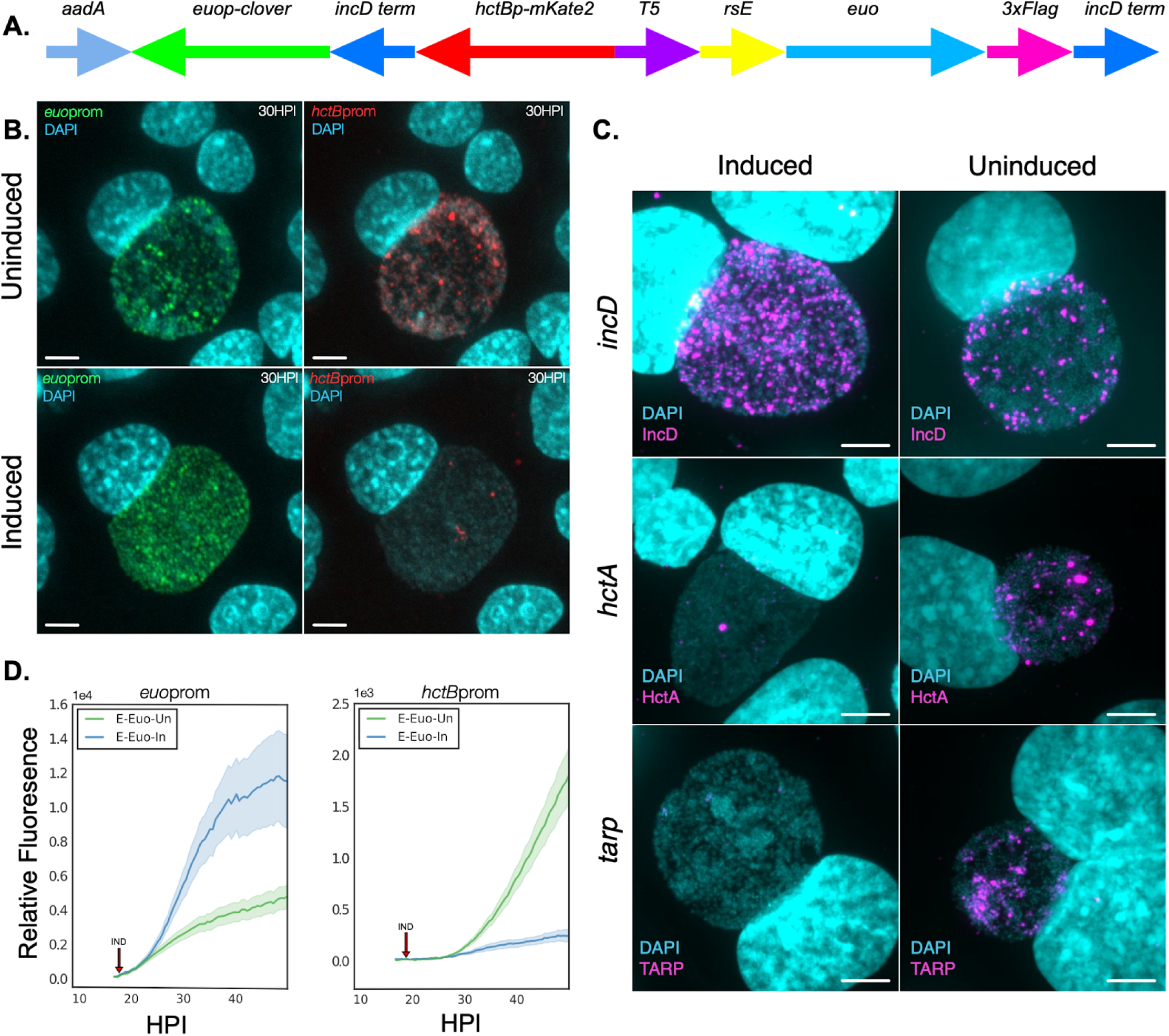
Characterization of the effects of ectopically expressed Euo on the chlamydial developmental cycle. A) Schematic of the E-Euo-BmEc construct which consisted of the E-Euo-FLAG construct expressed independently from the BmEc cassette featuring *euo*prom fused to Clover green fluorescent protein (*euo*p-clover), an incD terminator sequence (incD term), and *hctB*prom fused to mKate2 red fluorescent protein (*hctB*p-mKate2). B) Confocal micrographs of Cos-7 cells infected with L2-E-Euo-BmEc either untreated or treated with 0.5 mM Tph at 18 hpi and fixed at 30 hpi. The infected cells were imaged for DNA (DAPI, blue), *euo*prom signal (Clover, green), and *hctB*prom signal (mKate2, red). (scale bar = 15µm). C) Confocal micrographs of cells infected with L2-E-Euo-FLAG and stained using FISH for transcripts expressed in RBs (*incD*), IBs (*hctA*) and EBs (*tarp)*. Cells were Induced with Tph or left untreated (vehicle control) at 0 hpi and fixed at 24 hpi (scale bar = 15 µm). D) Live cell kinetics of *Ctr* ectopically expressing Euo. Cos-7 cells were infected with L2-E-Euo-BmEc, treated or not with 0.5 mM Tph at 18 hpi (red arrow) and imaged for 50 hours with automated live cell microscopy. Mean intensities are shown. Error cloud represents SEM. n > 20 inclusions per treatment.

To determine the effects of ectopic Euo expression on developmental kinetics, we utilized live cell microscopy and our dual promoter reporter system (2, 10, 11). Cells were infected with L2-E-Euo-BmEc and Euo expression was induced at 18hpi. The infected cultures were imaged every 30 minutes for 50 hours and relative fluorescence of red (*hctB*prom, EBs) and green (*euo*prom, RBs) channels measured for individual inclusions. The live cell expression kinetics of the two reporters showed a similar trend as the microscopy data, with an increase in *euo*prom and decrease in *hctB*prom expression in induced samples as compared to the uninduced cultures (Fig. 2D). Overall, these data suggest that ectopic expression of Euo arrested the chlamydial cells at an early stage of the developmental cycle.

### The Euo arrested cell population acts like IB cells and not RB cells

During the chlamydial developmental cycle, the RB cells replicate and ultimately produce IB cells through cell division. The IB cells exit the cell cycle and begin the process of developing into the infectious EB cell form (2). We used our previously published (2) agent-based model of the developmental cycle to simulate growth curves with cells arrested in the RB stage or arrested in the IB stage and compared that against our non-arrested simulation. If the *Chlamydia* were arrested as RBs our model predicted an extended exponential replication phase. However, when cells were simulated to be arrested in the IB state the replication kinetics was similar to the non-arrested simulation (Fig. 3A). We compared this simulation to induced and uninduced chlamydial growth curves generated using ddPCR to enumerate chromosomes over time. The growth curves were nearly identical for both the induced and uninduced samples, suggesting that ectopic Euo expression did not affect RB (replicating cells) amplification or IB/EB (non replicating cells) production (Fig. 3B). This data suggests that Euo is acting to block EB maturation from the committed IB cell form and not impacting the replicating RB cells.

**Figure 3:**
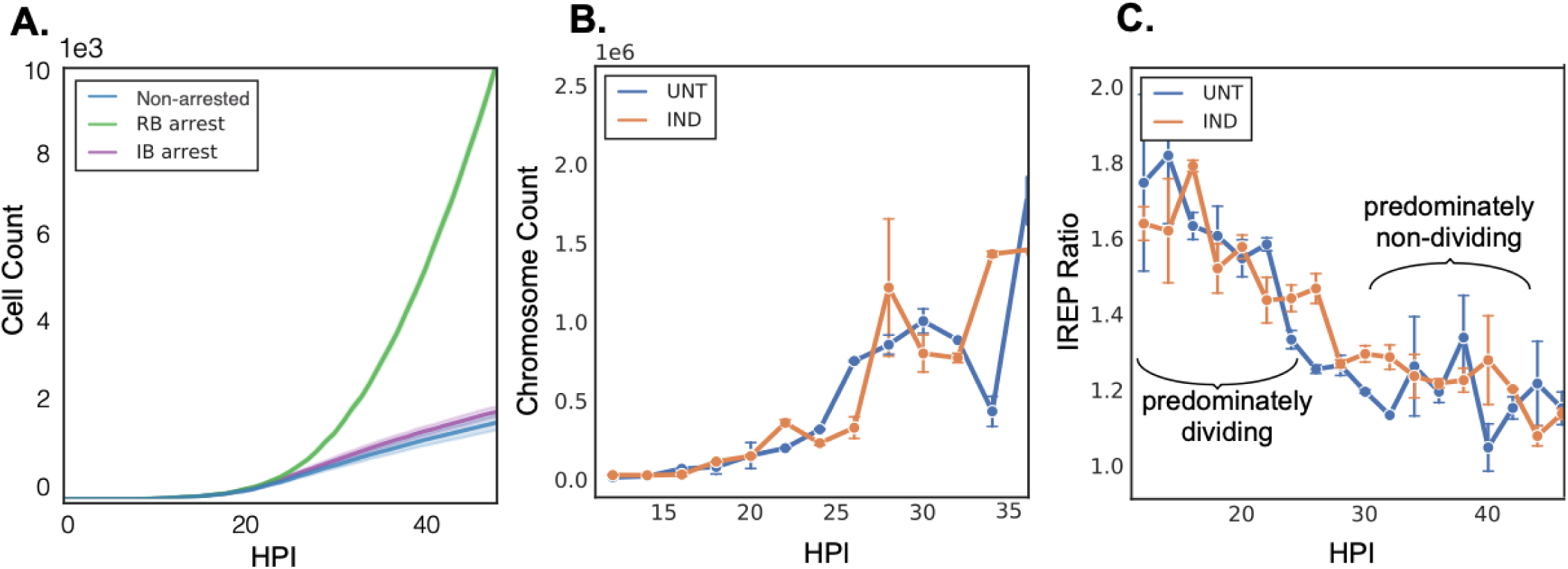
Chromosomal replication and modeling of the developmental cycle after Euo-FLAG ectopic expression. A) Computational modeling of *Ctr* development demonstrated population growth of different arrested cell forms. Simulations of both the non-arrested (blue) and the IB arrested (purple) chlamydial developmental cycle followed similar growth curves. Simulations of the RB arrested cycle (green) demonstrated logarithmic population growth. B) Chromosomal counts during growth of L2-E-Euo-FLAG untreated and induced. C) Index of Replication (iRep) of L2-E-Euo-FLAG untreated and induced populations. Induction occurred at infection and time points were taken every 2 hpi from 12-50 hpi. An iRep ratio >1.5 is considered a predominantly dividing population, an iRep ratio <1.5 is considered a predominantly non-dividing population. Error bar represents SEM. n > 50 droplets per treatment.

To further test whether ectopic Euo expression was affecting IB cell production and cycle exit, we calculated the replication index (iRep) of the infection over time. iRep is a measure of the ratio of the chromosomal origin vs the terminus resulting in an index that is proportional to replication rate (12, 13). An origin-to-terminus ratio close to 1 indicates the chromosome is not being replicated (ori and term concentrations are equal), while a ratio close to two indicates that most chromosomes are being replicated (13). We calculated the iRep index for induced and uninduced samples every 2 hours from 12 hpi to 50 hpi. The uninduced sample correlated well with our previously published data (12), i.e. the predominantly dividing cell phase (10-28 hpi) had a high index of replication (∼1.8 - ∼1.3) while the EB dominated phase (30-50 hpi) had a low iRep value (∼1.3 - ∼1.1) (Fig. 3C). In the Euo induced cell population the index tracked that of the uninduced; the index was the highest (∼1.8) early during infection and decreased over time until ∼30 hpi at which point the iREP remained low (∼1.1) for the remainder of the infection (Fig. 3C). These data suggest that ectopic Euo expression did not affect the formation of the early IB cell (cell cycle exit), nor did it affect RB replication.

### The Euo-dependent arrested developmental cycle is reversible

We next asked if the Euo arrested cell population could reenter the developmental cycle after Euo was no longer over expressed. We previously demonstrated that the developmental cycle follows a predictable kinetic gene expression pattern; the *hctA* promoter becomes active within a few hours of IB formation followed by *hctB*prom activation ∼8-10 hours later as the IB matures into the EB cell form (7). To determine the kinetics of recovery, cells were infected with L2-E-Euo-BmEc and induced Euo expression with Tph at the time of infection. We removed Tph at 24 hpi and measured the expression of the *hctB*prom reporter for 38 hours using live cell imaging (Fig. 4A). As reported in Fig. 2, ectopic expression of Euo repressed *hctB*prom activity as compared to the uninduced control (Fig. 4A, green and blue lines). When Tph was removed from the culture at 24 hpi, we observed a biphasic recovery of *hctB*prom activity; an immediate fast recovery that started ∼8 hours after washout followed by slower *hctB*prom production rate that matched the rate in the uninduced sample (Fig. 4A, purple and blue). We again used our agent based model of the developmental cycle to simulate recovery of *hctB*prom activity using the assumption that the arrested cell type were IBs (committed cell to EB formation). The simulations produced biphasic recovery kinetics that closely matched the measured data (Fig. 4A, purple and brown). Based on this simulation, the biphasic recovery was likely produced by an immediate recovery of the arrested early IB cell population followed by a return to a cell division dependent production of new IBs from the replicating RBs. These new IBs then matured into EBs. To test this prediction, we repeated the Tph washout experiment and inhibited cell division with penicillin (Pen). Cells were infected with L2-E-Euo-BmEc and either induced or not for ectopic Euo expression with Tph at the time of infection. Tph was removed at 24 hours and Pen was added. As seen previously, ectopic expression of Euo suppressed *hctB*prom activity compared to the uninduced (Fig. 4B, blue and green). Washout with the addition of Pen resulted in rapid *hct*Bprom recovery but the return to uninduced *hctB*prom activity kinetics (replication dependent, slow recovery) was blocked (Fig. 4B, purple). This data strongly supports the model that ectopic Euo expression arrested IBs at a very early stage but did not affect RB replication and production of new IBs.

**Figure 4:**
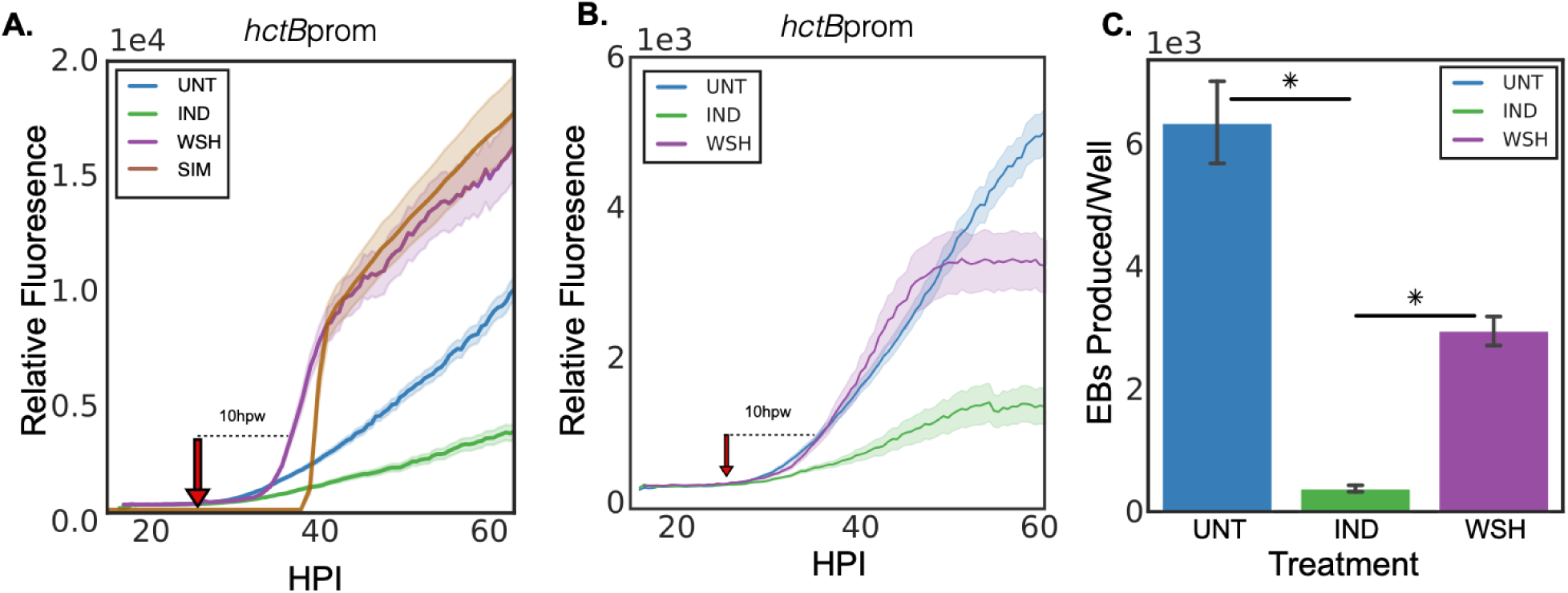
Recovery of the developmental cycle after Euo-FLAG induction and inducer washout. A) Live-cell imaging of Cos-7 monolayers infected with L2-E-Euo-BmEc induced or treated with vehicle control for Euo expression at infection. Cells were imaged starting at 10 hpi and Tph was removed at 24 hpi. For each experimental condition the fluorescent intensity of each promoter for individual inclusions was measured and the mean and SEM of >20 inclusions was plotted. Untreated control (blue line) showed the expected increase in *hctB*prom activity starting around 30 hpi. In the Tph induced sample (green line), *hctB*prom activity was repressed. After washout of the inducer (red arrow), *hctB*prom activity recovered in a biphasic manner (purple line); fast recovery starting ∼35 hpi and a return to wt *hctB*prom kinetics ∼40 hpi. Computational simulation of the developmental cycle using the assumption that the arrested cells are in the IB state generated similar biphasic recovery kinetics (brown line). B) Live-cell imaging of chlamydial inclusions from Cos-7 monolayers infected with L2-E-Euo-BmEc treated with Pen after Tph washout. Cells were infected and either induced to express Euo or treated with vehicle only at infection. Inclusions were imaged starting at 10 hpi. In the induced samples *hctB*prom activity was inhibited (green line) while the uninduced *hctB*prom activity demonstrated the expected linear increase from ∼30 hpi until the end of the experiment (blue line). Recovery in the washout experiment (removal of Tph and addition of Pen at 24 hpi, red arrow) showed only the early fast phase of *hctB*prom recovery and an inhibition of the later slow phase of recovery (purple line). Error cloud for fluorescent reporters represents SEM. n > 20 inclusions per treatment. C) IFU assay of recovery. Cos-7 monolayers were infected at an MOI ∼1 with L2-E-Euo-BmEc. Euo expression was induced at 0 hpi and EBs were harvested at 48 hpi. The harvested EBs were used to reinfect a monolayer and inclusions were enumerated. Asterisks = p < 0.01.

The kinetic recovery data suggested that ectopic expression of Euo arrested chlamydial IB cells in an early IB state, and that when induction was reversed these IBs reentered the developmental cycle ultimately resulting in *hctB*prom activation. To verify that recovery resulted in the production of EBs, we used confocal microscopy to visualize the chlamydial cells and measured the activity of the *euo*prom and *hctB*prom in individual *Chlamydia*. Cos-7 cells were infected with L2-E-Euo-BmEc and induced with Tph at infection. Tph was washed out at 24 hpi and the infected cells were fixed and stained for FLAG expression and imaged using confocal microscopy at 35 and 38 hpi which corresponded to 11 hours and 14 hours post washout, respectively (Fig. S2A). Confocal images from 38 hpi show that in the Tph induced cultures the chlamydial cells expressed clover from *euo*prom and very few cells expressed mKate2 from *hctB*prom. After washout very little ectopic Euo expression was detected (FLAG staining) and at 38 hpi there was a significant increase in the number of *hctB*prom+ cells and these appeared to be EB like (Fig. S2A). We quantified the expression of *euo*prom and *hctB*prom in individual chlamydial cells at 35 and 38 hpi and plotted the fluorescence (Fig. S2B). Using Trackmate and FIJI (14) we identified each *euo*prom+ cell (green dots) and each *hctB*prom+ cell (red dots) and measured the fluorescence of each cell in both channels. We plotted the *hctB*prom intensity against *euo*prom intensity at 24 hpi (induced), 35 hpi (induced), 38 hpi (induced) and 35 and 38 hpi after washout of Tph at 24 hpi (Fig S2B). At 24 hpi in the induced population the majority of the cells were only *euo*prom+. This was true at 35 and 38 hpi when Tph was still present (Fig S2B). Upon washout the number of *hctB*prom+ cells increased and these cells had little to no *euo*prom signal (Fig. S2B). There was also an increase in *hctB*prom signal in a population of the *euo*prom+ cells which are likely late IBs that are in the process of transitioning from *euo*prom+ to *hctB*prom+ (Fig. S2B). This data suggests that the arrested population recovered and matured into EBs. To directly test whether the recovered population was infectious, we performed an IFU reinfection assay. Monolayers were infected with the L2-E-Euo-FLAG chlamydial strain and induced at 0 hpi. We then washed out Tph at 24 hpi and harvested the chlamydial cells at 48 hpi. These cells were tested for infectivity using an inclusion forming replating assay (Fig. 4C). The Tph treated population had very few infectious EBs while washout resulted in a significant increase in infectious EBs (Fig. 4C).

In addition to the IFU assay we also used live cell microscopy to visualize the formation of re-infection plaques after Tph washout. Cos-7 cells were infected with L2-E-Euo-BmEc and induced with Tph at infection. Tph was washed out at 24 hpi and the infected monolayer was imaged for an additional 40 hours. The Tph treated sample without washout had inclusions with very little *hctB*prom signal and no re-infection plaques were formed during the imaging period. However, in the washout experiment there was a dramatic increase in *hctB*prom signal in a subset of inclusions. These inclusions increased in size and after lysis created plaques of newly infected neighboring cells (Movie S1 and S2). We also noticed that upon Tph washout there was an almost immediate increase in inclusion lysis. In the washout samples, ∼35% of the inclusions lysed within three hours of Tph washout (Fig S3). This was much higher than for the non-washed out inclusions (∼12%) while there was essentially no lysis during this time frame (∼1%) for the untreated samples. The increase in lysis for the washout experiment likely explains the partial recovery of infectious progeny observed in Fig 4C.

### The Euo arrested cells appear stalled in cell division

Our data indicated that ectopic Euo expression arrested chlamydial cells in an early IB like state. Like true IBs and EBs these cells are out of the cell cycle and not replicating, but are not expressing IB or EB genes. *Chlamydia* do not encode the *ftsZ* gene and instead construct a peptidoglycan ring that functions in cytokinesis (15, 16). Therefore, to determine the cell division status of the Euo arrested cell form, we used bioorthogonal click chemistry to label the peptidoglycan ring of uninduced, induced and induced+washout samples. Cos-7 cells were infected with L2-E-Euo-BmEc and induced with Tph or treated with vehicle at the time of infection. Ethynyl-D-alanyl-D-alanine (EDA-DA, Thermo Fisher Scientific) was added at 20 hpi to label the peptidoglycan (15). The cells were fixed and the peptidoglycan was stained using click chemistry at 24 hpi (17). In the uninduced cells there were obvious EBs (small condensed DAPI stained cells) and RBs (large dispersed DAPI stained *euo*prom+ cells). The RB cells in this population had obvious large rings around some RBs and smaller rings between septating dividing RBs (Fig. 5A, 24hpi UNT). The small EB cells did not have peptidoglycan rings. In the population induced for Euo expression, there are essentially no EB cells, only *euo*prom+ cells and, like the uninduced population, the *euo*prom+ cells had visible peptidoglycan rings associated with them. There were cells that appeared to be actively dividing with small rings separating RB like cells (Fig. 5A, 24hpi IND). However, there was an increase in a population of cells that appeared stalled in cell division with a full sized ring midway around slightly elongated cells (Fig. 5A, 24hpi IND). We investigated the ability of the induced cells to recover when Euo was no longer overexpressed. For this experiment Tph was washed out at 24 hpi and the cells were fixed four hours later. After washout, the chlamydial cells appeared to recover and looked similar to that observed in uninduced samples (Fig. 5A, 24hpi IND +4 hpw). There were again small cells with dense DAPI staining of which most did not have peptidoglycan rings while the rings of the *euo*prom+ population had more intermediate cell division forms with rings of various sizes visible between dividing cells.

**Figure 5:**
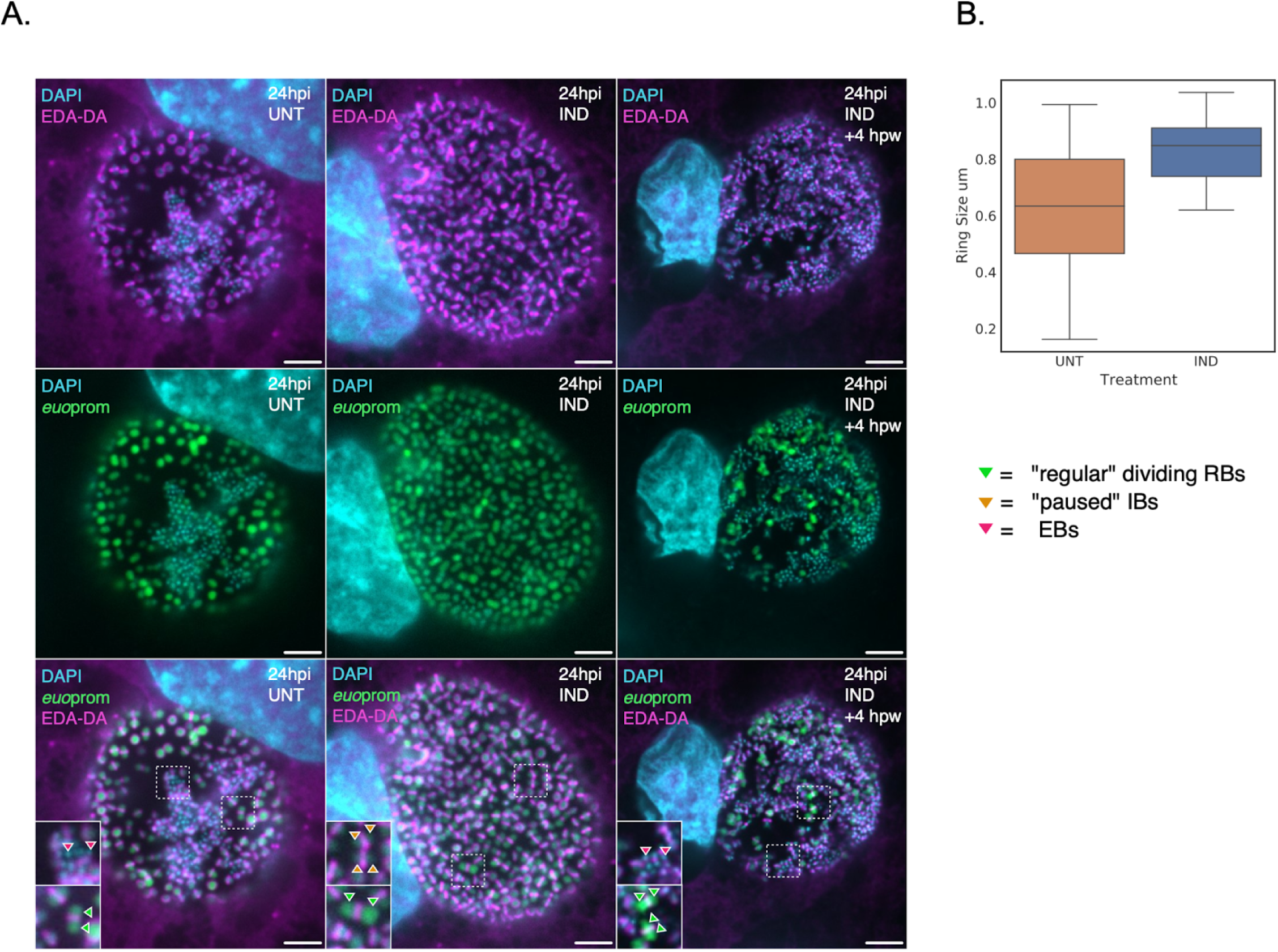
Visualization of the peptidoglycan division ring after Euo induction and washout. A) Cells were infected with L2-E-Euo-BmeC and either induced for Euo expression with Tph or vehicle only at infection. At 20 hpi the infected cells were fed ethynyl-D-alanyl-D-alanine (EDA-DA) and fixed at 24 hpi. EDA-DA was incorporated into the peptidoglycan of *Ctr* and was visualized using a fluorescent azide and Click chemistry. Confocal single slice images of *Ctr* with the peptidoglycan division rings labeled with Alexa 640 (megenta), DNA labeled with DAPI (blue) and visualization of *euo*prom activity (green). The uninduced sample showed peptidoglycan rings around green RB cells at different stages of cell division (inset) and EB cells with no visible peptidoglycan rings (inset). The induced population had very few EBs and an increase in *euo*prom+ cells with arrested division rings (inset) in addition to RB division intermediates (inset). Tph was removed and *Ctr* was allowed to recover for four hours before fixation and staining. In the recovered population there were again peptidoglycan rings around green RB cells at different stages of cell division (inset) and EB cells with no visible rings (insets). Scale bar = 10µm. B) Quantification of peptidoglycan ring size. The diameter of the rings in 20 Inclusions were measured and plotted. The ring sizes were highly variable in the uninduced samples with full sized rings, intermediate sized rings and small rings present between septating RBs. In the Tph treated population the rings were almost all large, fully encircling the *euo*prom+ cells. The central line represents the median, while the top and bottom of the box represent 75th and 25th percentiles respectively.

The dramatic increase in stalled full sized rings was quantified by measuring ring diameters using FIJI. In the uninduced cell population there were dividing RBs, IBs transitioning to EBs and EB cell forms. The RB cells had peptidoglycan division rings of various sizes (Fig. 5B), both fully encircling the RB cells or small rings separating septating/dividing cells. The Euo expressing population had an overall dramatic increase in ring sizes within the population (Fig. 5B), as dividing RB forms with small rings were still present but the majority of the chlamydial cells had rings that fully encircled the cells. These data suggest that the Euo arrested cell form is trapped in an intermediate stage. The cells have fully replicated their chromosomes as indicated by the iRep data but have not yet fully divided into two daughter cells. These cell division arrested cells were able to complete division and form EBs after the inducer was removed.

### Arrest and reentry of the developmental cycle by expression of Euo was similar for both the synthetic T5 promoter and the native *euo* promoter

Our data showed that ectopic expression of Euo using the *E. coli* synthetic T5 promoter resulted in arrest of the developmental cycle at an early stage of IB/EB development. Upon washout of the Tph inducer the cycle resumed. To determine if this same behavior would be observed using the native *euo* promoter to drive ectopic expression of *euo* mRNA (induced for protein expression by Thp), i.e. in the the correct cell form at the correct time, we constructed a translationally regulated expression construct using the *euo* native promoter, a riboJ insulator and the riboE Tph sensitive riboswitch as described previously (10). We replaced the T5-riboE promoter driving Euo-FLAG expression with this regulatable native promoter creating Nativeprom-Euo-3xFLAG-*hctB*prom-NeonGreen (*euo*nprom-E-Euo-Bng) and the strain L2-*euo*nprom-E-Euo-Bng (Fig. 6A).

**Figure 6:**
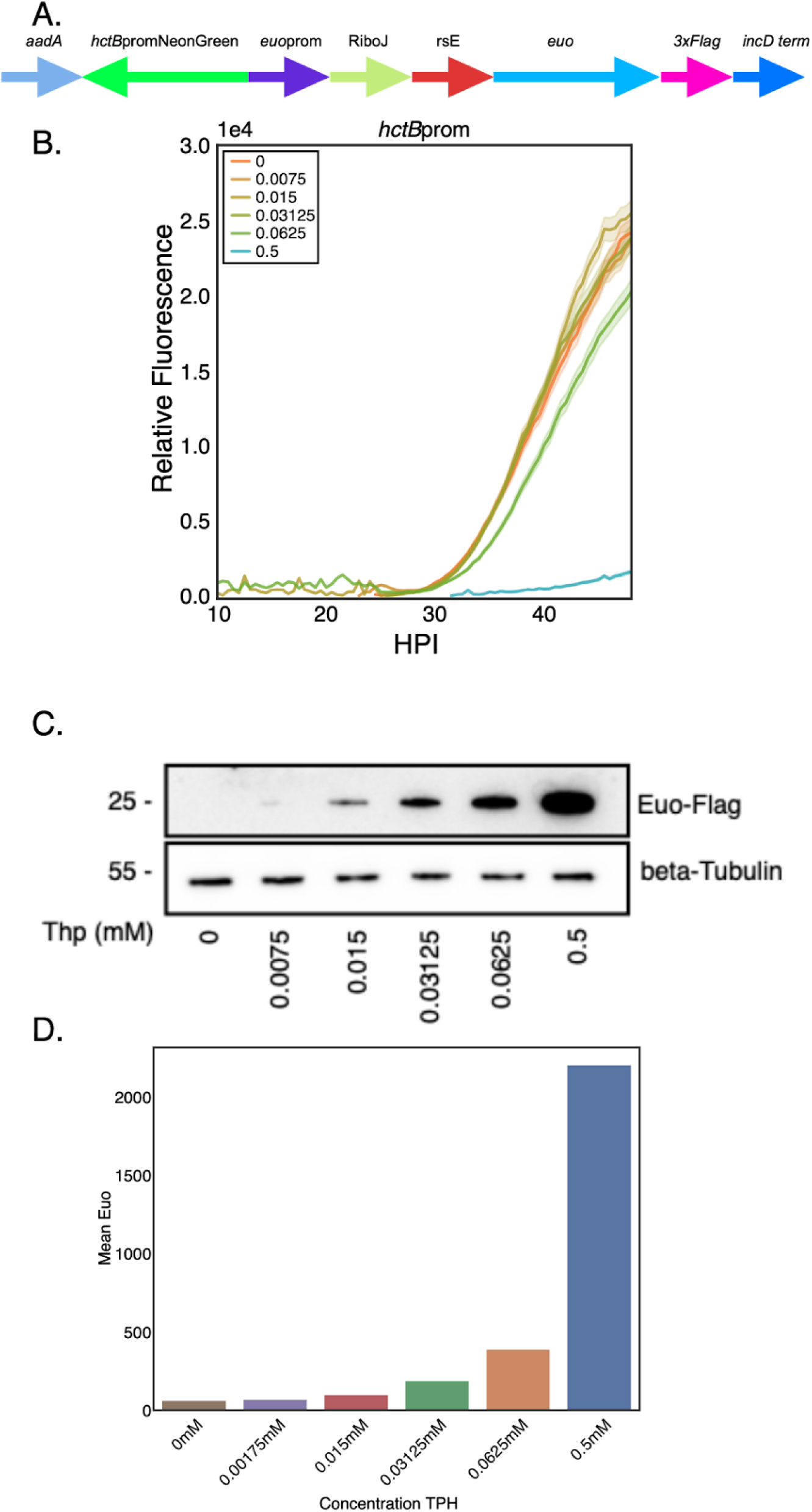
Euo levels control developmental cycle arrest. A) Schematic of e*uo*prom-E-Euo-Bng construct which consisted of the native *euo*prom driving the expression of Euo-FLAG independently from the *hctB*prom-NeonGreen promoter reporter cassette. B) Fluorescent live-cell microscopy of L2-e*uon*prom-E-Euo-Bng infected cells induced with various concentrations of Tph resulted in a biphasic repression of *hctB*prom activity. Cells were infected with L2-*euo*nprom-E-Euo-Bng and induced with 0.5. 0.0625, 0.03125, 0.015, 0.00175 and 0 mM Tph at 0 hpi. Samples were imaged using live cell imaging and the fluorescence signal for individual inclusions was determined for > 20 inclusions from 10 hpi until 50 hpi. Error cloud for the fluorescent reporter represents SEM. C) Western blotting of the induced samples at 24 hpi using an anti-FLAG antibody. D) Densitometry quantification of western blots.

We measured the effects of Euo expression on the kinetics of the developmental cycle using live cell imaging. Cells were infected with L2-*euo*nprom-E-Euo-Bng and treated with 0.5, 0.06, 0.03, 0.015, or 0.0075 mM Tph at infection and imaged for 48 hours capturing images every 30 minutes. At the highest Tph dose we observed developmental cycle arrest similar to that observed with the L2-E-Euo-BmEc strain (Fig. 6A compared to Fig. 2D). However, at the other doses we saw very little effect on the developmental cycle kinetics as measured by *hctB*prom activity (Fig 6B). We measured the expression levels of Euo-FLAG by western blotting and found that a level of Tph as low as 0.015 resulted in detectable levels of expression (Fig. 6C and D). We repeated the kinetic experiment with more closely spaced Tph dilutions and observed a non-linear effect on the developmental cycle. The 0.5 mM, 0.25 mM and 0.125 mM concentrations significantly reduced *hctB*prom activity while the 0.0625 mM, 0.031 mM, 0.015mM and 0mM concentrations had limited effects on the progression of the developmental cycle (Fig S4). These data demonstrated that arrest of the developmental cycle by Euo overexpression occurs for both the native *euo* promoter as well as the synthetic *E. coli* sigma70 promoter. The data also suggested that the resumption of the developmental cycle is controlled by a feed-forward loop; as Euo levels drop below a threshold, the cycle can move forward.

### Euo auto regulates

Overexpression of Euo from a plasmid using either the T5 promoter or the native *euo* promoter resulted in reversible arrest of the developmental cycle. This suggested that Euo levels control forward progression of the cycle with Euo acting as a switch for IB/EB maturation. To further investigate the role of Euo on Euo regulation we determined the effects of Euo ectopic expression on Euo protein levels. Cells were infected with either L2-E-Euo-BmEc or L2-*euo*nprom-E-Euo-Bng and Euo expression was induced at infection. The infected cells were either fixed at 24 hpi or Tph was washed out and the *Chlamydia* were allowed to recover for two and four hours before fixation. The cells were stained for Euo protein levels using an anti-FLAG antibody. Confocal microscopy revealed that Euo-FLAG was abundantly expressed in arrested cells at 24 hpi. This was true for cells infected with either strain (Fig. 7A and C, Induced). The FLAG signal was dramatically reduced four hours after washout for both strains (Fig. 7A and C, 24hpi + 4 hpw). We measured the fluorescent intensity of the FLAG staining in five Inclusions using summed z stacks and FIJI at 24 hpi (before washout) and at two (26 hpi) and four (28 hpi) hours after washout. Washout of Tph resulted in a dramatic reduction in FLAG staining as soon as two hours post washout for both L2-*euo*nprom-E-Euo-Bng and L2-E-Euo-BmEc (Fig. 7B and D).

**Figure 7:**
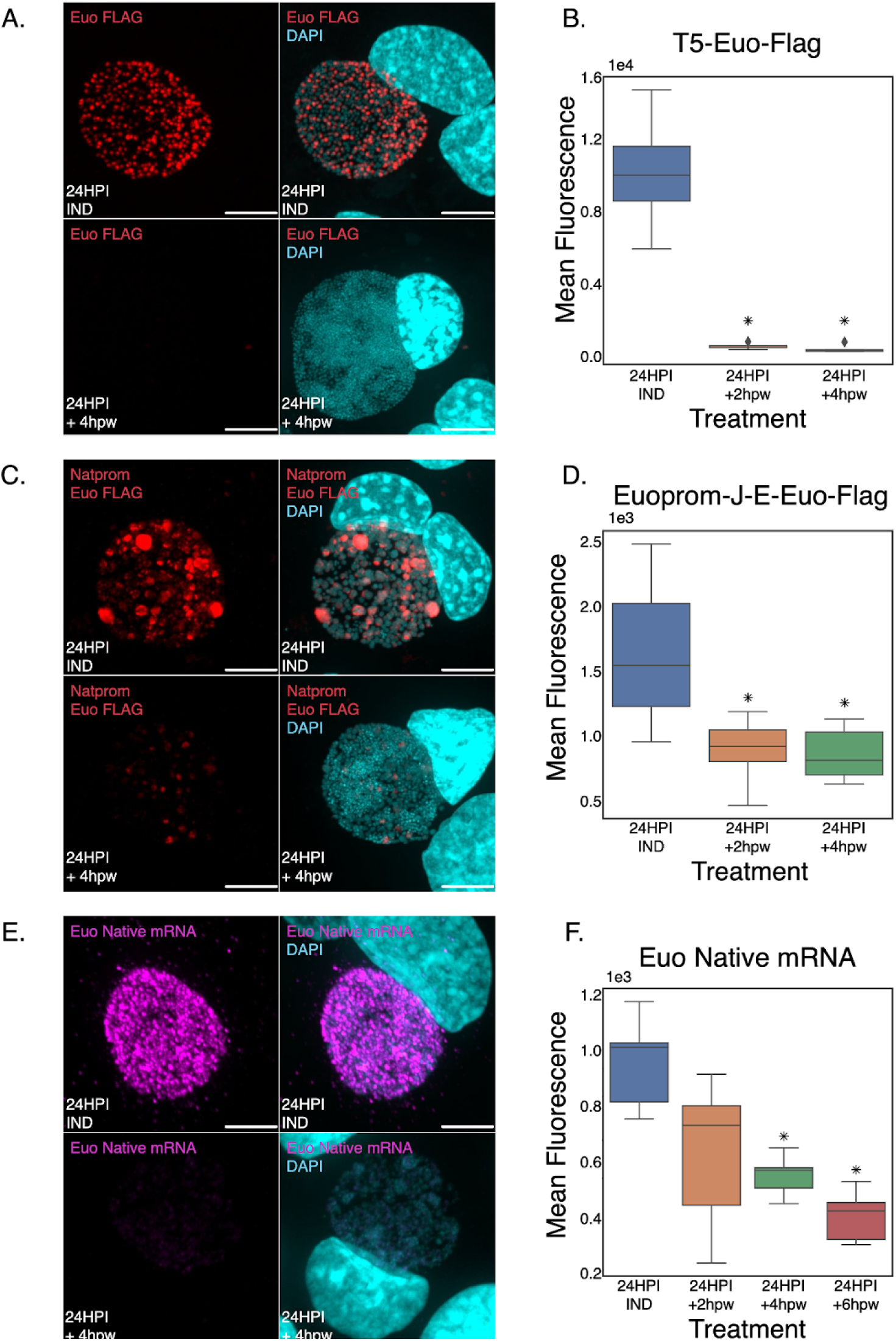
Levels of ectopically expressed Euo-FLAG and chromosomal *euo* mRNA are reduced upon inducer washout. A) Confocal micrographs of L2-Euo-FLAG infected cells induced at infection with Tph, stained for the FLAG tag (red) and DAPI (blue) at 24 hpi and four hours after inducer washout (+4hpw). (scale bar = 15µm). B) Quantification of mean Euo-FLAG fluorescence of L2-E-Euo-BmeC infected cells induced at infection with Tph. Time points were taken at 24 hpi (pre-washout), 26 hpi (+2hpw), and 28 hpi (+4hpw). Fluorescence of 8 inclusions measured per treatment. Asterisks = p < 0.01 as compared to 24 hpi. C) Confocal micrographs of L2-*euon*prom-E-Euo-Bng infected cells induced at infection with Tph, stained for the FLAG tag (red) and DAPI (blue) at 24 hpi and four hours after inducer washout (+4hpw). (scale bar = 15µm). D) Quantification of mean Euo-FLAG fluorescence of L2-*euo*nprom-E-Euo-Bng infected cells induced at infection with Tph. Time points were taken at 24 hpi (pre-washout), 26 hpi (+2hpw), and 28 hpi (+4hpw). Fluorescence of 8 inclusions was measured per treatment. Asterisks = p < 0.01 as compared to 24 hpi. E) Confocal micrographs L2-*euo*nprom-E-Euo-Bng infected cells induced at infection with Tph, stained for the chromosomal transcript (magenta) and DAPI (blue) at 24 hpi and four hours after inducer washout (+4hpw). (scale bar = 15µm). F) Quantification of mean FISH fluorescence of the chromosomal *euo* transcript from the L2-euoprom-E-Euo-Bng infected cells induced with Tph. Time points were taken at 24 hpi (pre-washout), 26 hpi (+2hpw), 28 hpi (+4hpw) and 30 hpi (+6hpw). Fluorescence of 8 inclusions was measured per treatment. Asterisks = p < 0.01 as compared to 24 hpi.

We next saught to determine the fate of chromosomally encoded *euo* mRNA expression. The *euo*nprom-E-Euo-Bng plasmid contained a synonymous codon substituted *euo* gene so that native *euo* mRNA could be differentiated from ectopically expressed mRNA. Cos-7 cells were infected with L2-*euo*nprom-E-Euo-Bng, induced with Tph and fixed and stained for the native *euo* mRNA at 24 hpi and four hours post Tph washout. At 24 hpi the arrested cells demonstrated high native *euo* mRNA FISH staining (Fig. 7E, Induced). After washout and allowing the arrested *Chlamydia* to reenter the developmental cycle for 4 hours, FISH staining revealed that the native *euo* mRNA was dramatically reduced in the recovered cell population (Fig. 7E, 24hpi + 4 hpw). We measured the fluorescent intensity of the FISH signal in five inclusions using summed z stacks and FIJI at 24 hpi (before washout) and at two, four and six hours after washout. Similar to the ectopically expressed Euo protein, the native *euo* mRNA decreased rapidly after washout (Fig. 7B). Taken together, these data suggest that Euo acts in a feed-forward loop, i.e. when Euo levels are high Euo is expressed, but when Euo levels drop Euo expression becomes repressed.

## Discussion

Bacteria of the *Chlamydiaceae* family are all obligate intracellular parasites of vertebrate cells (18). *Chlamydia* have no environmental reservoirs and must actively invade and replicate in host cells to survive. A central goal for all bacteria is to increase cell numbers and disseminate to new environments. To accomplish this, *Chlamydia* has evolved a complex multi-phenotypic cell type developmental cycle. The chlamydial developmental cycle produces highly adapted phenotypic cell forms specialized for either dissemination or replication (19, 20). The dissemination-specialized form is the chlamydial EB, a non-replicating cell form with reduced transcriptional and translational activity (21, 22). The EB is characterized by its small size (∼0.2 µm diameter) and highly condensed nucleoid (20). This form mediates host cell attachment and invasion (9). The RB cell form is larger (∼1 µm diameter) and replicates inside the host cell. Replication takes place in a membrane bound compartment heavily modified by *Chlamydia* to create a replication niche termed the chlamydial inclusion (23–25). The process of producing EBs appears to involve asymmetric division by the RB cell producing an intermediate cell form, the IB (2). The chlamydial IB is the committed cell form that directly develops into the infectious EB. Chlamydial replication amplifies RB numbers and produces EBs that can then disseminate the infection to new host cells for subsequent rounds of infection (2). The regulatory circuits that control this complex developmental cycle are incompletely defined.

*Chlamydia* encodes a DNA binding protein Euo that is expressed early after cell entry and only in the RB cell form (2, 6). Euo is a helix-loop-helix DNA binding protein that has been shown to act as a repressor of chlamydial late genes and a potential activator of mid cycle genes (4). Our studies here show that ectopic expression of Euo resulted in an arrest of the developmental cycle. The arrested cells have a gene expression profile similar to RBs, do not express IB or EB genes and appear to be out of the cell cycle and not replicating. The Euo arrested cells reentered the cycle when the inducer was removed. Interestingly, recovery was biphasic, with a fast recovery component followed by a return to the developmental kinetics of the uninduced controls. Use of our agent-based computational model of the developmental cycle (2) suggested that biphasic recovery was best explained by the accumulated arrested cells reentering the developmental cycle as IBs and synchronously recovering while the second slower recovery phase returned to replication dependent production of IBs. This interpretation was supported by the observation that inhibiting cell division blocked slow recovery but not fast recovery.

For the arrest and recovery experiments, Euo was expressed using the synthetic T5 promoter which is a strong constitutive sigma70-dependent promoter from *E. coli* (26). We substituted the T5 promoter with the native *euo* promoter but added riboswitch translational control (10) and the kinetics of developmental cycle arrest and recovery were the same. The data also showed that arrest was expression level dependent, however this did not follow a linear response. Instead, Euo-induced arrest appears to be dependent on a threshold of Euo expression. Recovery from arrest was relatively fast suggesting that the arrested cell is primed to turn off Euo expression and move forward in the cycle. The data show that within two hours of removing the Tph inducer, Euo-FLAG levels are almost undetectable. This was true for both T5 promoter expressed Euo-FLAG as well as for the native *euo* promoter regulated Euo-FLAG. Importantly, the loss of Euo-FLAG also resulted in the repression of chromosomally encoded *euo* as shown by FISH staining for the native transcript. These data suggest Euo is acting in a feed forward loop that is involved in shifting chlamydial gene expression from RB like to IB/EB like expression patterns. This data also suggests that Euo represses its own repressor; as soon as Euo levels drop *euo* gene expression is suppressed. It’s unclear how Euo protein levels are reduced in the IB cell form; this could be due to asymmetric partitioning during IB formation or, more likely, due to increased turnover in the early IB.

Our iRep data showed that the Euo arrested cell form, like the IB and EB, is out of the cell cycle and not replicating its chromosome. Intriguingly, this arrested form is not completely out of the division cycle as they had large peptidoglycan division rings encircling them. This observation leads to a compelling model of the developmental cycle. In our previous published model of the cycle, IBs are created from asymmetric division of the mature RB_E_ (IB producing RBs) followed by maturation of the IB directly to the EB without further cell division (2). However, our data presented here suggests that the IB cell may have a more complicated role in EB formation. The Euo arrested cell data strengthens our model that one of the RB_E_ daughter cells becomes an IB and exits the cell cycle. However, this exit appears to be delayed by one cell division. The daughter cell destined to become the IB likely completes one final round of chromosome replication without significant increase in cell size (Fig. 8). This early IB then divides one last time resulting in two daughter cells that are reduced in size, each containing one fully replicated chromosome. This model is additionally supported by the observations from Lee et al. that showed that the average size of the dividing RB like cell form decreases during infection leading to EB formation (27). Together these observations fit a model wherein RBs divide followed by cell growth. A subset of these RB cells (RB_E_s) divide asymmetrically producing an IB cell that does not grow in size but finishes one round of chromosome replication before a final division producing two smaller daughter cells. These two small daughter cells then transition into the EB form. This process would guarantee that each EB contains one fully replicated chromosome. We hypothesize that ectopic expression of Euo resulted in a cell form that was arrested at this early IB stage. This arrested cell is out of the cell cycle, contains two complete chromosomes and is surrounded by a fully formed septation ring. This cell state would explain the observed biphasic recovery upon inducer washout. After inducer washout the arrested early IB immediately divides one last time and initiates EB maturation (fast recovery), followed by a return to RB_E_ division dependent IB production (slow recovery).

**Figure 8:**
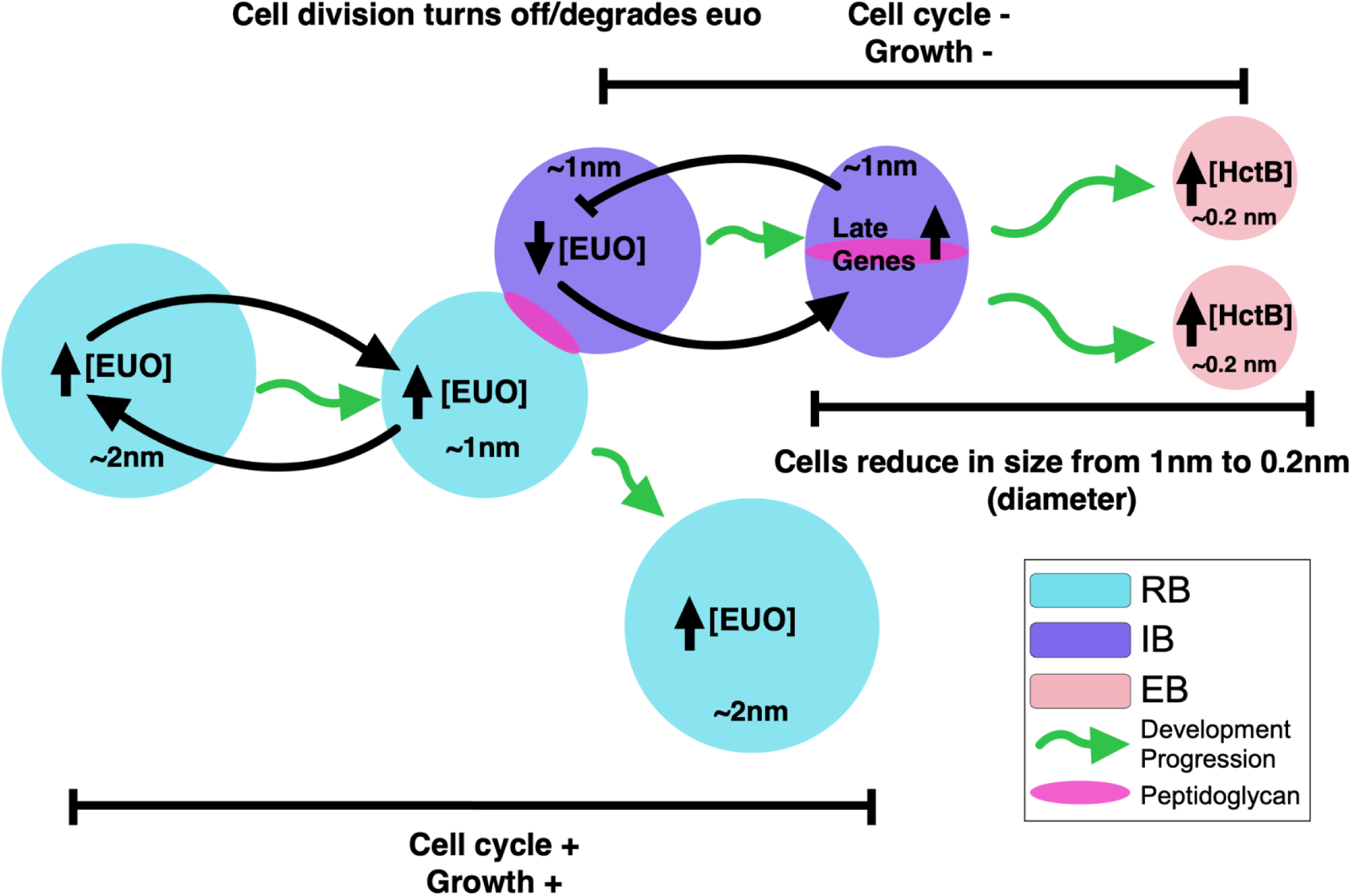
Model of the role of Euo in chlamydial cell form development. In this model, Euo is highly expressed in RB’s repressing IB and EB genes and upregulating itself via a feed forward loop. After asymmetric cell division one daughter cell becomes committed to EB formation and is considered an IB. In the IB form Euo levels drop leading to *euo* repression acting as a switch, repressing RB genes and activating the IB genes (*hctA*) and eventually the EB genes (*hctB*). The IB cell finishes one more round of chromosomal replication before dividing into two cells that become the EBs, each with a single fully replicated chromosome. This last division completes without cellular growth leading to an overall cell size reduction. Green arrows = developmental progression.

These data suggest an overall model of the developmental cycle that begins with EB entry and germination resulting in the expression of Euo. This is followed by RB_R_ replication increasing RB numbers, and then RB_E_ replication, producing IB forms. In the very early IB, Euo levels are reduced through an unknown mechanism leading to repression of Euo expression (Fig. 8). Falling levels of Euo act as a feed forward loop leading to derepression of IB/EB genes. The early IB cell form then divides one last time to produce two cells that further mature to EBs (Fig. 8). This overall model suggests novel mechanisms to produce the unusual EB cell which is out of the cell cycle, contains one fully replicated chromosome and is dramatically reduced in cell size. Additionally, these data suggest that Euo expression is a key switch in IB to EB maturation but is not involved in the committed step to IB formation.

## Materials and Methods

### Cell Culture

Cos-7 cells obtained from ATCC were maintained using RPMI 1640 supplemented with 10% fetalplex and treated with 10mg/mL gentamicin to prevent contamination. Cultures were grown at 37 degrees Celsius with 5% CO_2_. All *C. trachomatis* L2-bu434 (L2) infections were carried out in Cos-7 cells. EBs isolated from infectious cultures were harvested and purified utilizing centrifugation over a 30% MD-76R density gradient. Purified EBs were stored in sucrose-phosphate-glutamate buffer (SPG) (10mM sodium phosphate [8mM K_2_HPO_4_, 2mM KH_2_PO_4_], 220mM sucrose, and 0.50 mM L-glutamic acid, ph 7.4) at -80 degrees Celsius.

### Reporter Plasmids

All reporter plasmids were made using the p2TK2SW2 backbone (28). Promoters from *Chlamydia* were amplified using *C. trachomatis* L2 genomic DNA using the primers in Table S1. All promoter sequences started ∼100bp from the predicted transcription start site and included the non translated region as well as the first 30bp (or first 10 amino acids) of the ORF for the specified gene. Fluorescent reporters (Clover/mKate2) were ordered from Integrated DNA Technologies (IDT) as gene blocks and inserted in frame with the promoter/+30 gene. The *aadA* gene for spectinomycin resistance was added from pBam4. Each ORF ended with an incD terminator. This resulted in the plasmids p2TK2-E-Euo-3xFLAG (E-Euo-FLAG), and p2TK2-E-Euo-3xFLAG_hctBprom-mkate2_euoprom-mClover (E-Euo-BmEc). To make the *euo* native promoter construct, *euo*nativeprom-riboJ-E-Euo (codon substituted)-3xFlag_*hctB-*neongreen (euonprom-E-Euo-Bng) the T5 promoter in front of the E riboswitch was replaced with the euo native promoter and the ribo J insulator (10). Additionally, the euo open reading frame was replaced with a codon substituted euo gene. The majority of the codons were substituted with alternate codons so that the ectopically expressed *euo* could be distinguished from native *euo* mRNA using FISH. The codon modified and native Euo genes were tested for homology using Blast and no homology was reported (29). This construct was ordered from IDT as a double stranded DNA gBlock (EuoProm-J-E-Euo-(codon substituted)-Flag_gblock). The dual color reporter cassette BmEc was replaced with the single color reporter *hctB*prom-neongreen. This reporter cassette was also purchased from IDT as a double stranded gBlock (hctBneongreen-synterm_gblock).

### Chlamydial Transformation

*C. trachomatis* transformation was performed as previously described (10, 30) using 500ng/ul spectinomycin for selection. Clonal populations were isolated by titration of the infection followed by inclusion isolation with a micromanipulator. The plasmids from the chlamydial transformants were verified by sequencing.

### Infections

Cos-7 cell monolayers were incubated with infectious EBs in Hanks Balanced Salt Solution (HBSS) (Gibco) for 15 min at 37 degrees Celsius. The inoculum was then removed and the host cells were washed with pre-warmed HBSS. After washing the HBSS was replaced with fresh RPMI 1640 containing 10% fetal bovine serum, 10 μg/ml gentamicin, 1 μg/ml cycloheximide, and 1 mg/ml heparin to ensure synchronization of infection and to lower reinfection in long experiments.

### Transmission Electron Microscopy

For analysis of the structure of *Ctr* upon ectopic protein expression, cell monolayers were infected with the indicated strain at an moi of 0.5 and induced with 0.5mM Tph at 15 hpi. Infected cells were released from the plate with Trypsin-EDTA at 30 hpi, rinsed with 1xPBS and the pellet was fixed with EM fixative (2%PFA, 2% Glutaraldehyde, 0.1M Phosphate Buffer, pH 7.2) overnight at 4°C. Fixed pellets were rinsed and dehydrated before embedding with Spurr’s resin and cross sectioned with an ultramicrotome (Riechert Ultracut R; Leica). Ultra-thin sections were placed on formvar coated slot grids and stained with uranyl acetate and Reynolds lead citrate. TEM imaging was conducted with a Tecnai G2 transmission electron microscope (FEI Company; Hillsboro, OR).

### Live Cell Microscopy

Cos-7 monolayers were grown on glass bottom 6-well plates and temperature and CO_2_ was maintained using an OKOtouch stage incubator. Infections were treated at 0 hpi with 0.5mM Tph unless otherwise stated. A Nikon Eclipse TE300 inverted microscope was used for live cell imaging using a 20x, 0.4-numeric-aperture objective. Fluorescent protein excitation was achieved using a ScopeLED lamp at 470 nm and 595 nm along with BrightLine bandpass filters at 514/30 nm and 590/20 nm. DIC was used for focusing. Image acquisition was achieved using a Andor Zyla sCMOS camera. Micro-Manager software was used to image every 30 min. Imaging started at 8 - 10 hpi unless otherwise stated until 80 hpi to fully visualize the developmental cycle. Induction washouts were achieved using a peristaltic pump system controlled by python scripts. Experiment data analysis utilized matplotlib, pandas, and seaborn using custom python notebooks to determine and utilized trackmate software to measure fluorescence intensities (2, 11).

### Confocal Microscopy

infected monolayers were fixed overnight in 2% paraformaldehyde (PFA) at 4 degrees Celsius. They were washed the next day with phosphate buffered saline (PBS). Staining for FLAG was achieved using monoclonal anti-FLAG M2 antibody (1:1,000, Sigma, Thermo Scientific™). Alexa 647 Goat-anti Mouse IgG-HRP secondary antibody (Invitrogen™) was used for fluorescent tagging of FLAG. Coverslips were mounted using MOWIOL mounting solution (100 mg/mL MOWIOL® 4–88, 25% glycerol, 0.1 M Tris pH 8.5). Confocal images were acquired using a Nikon CrestOptics X-Light confocal coupled with Nikon Elements imaging software. Images were taken using a 100x oil-immersion objective.

### Replating Assay

*Ctr* were isolated by scraping infected monolayers and pelleting at 18213 rcf for 30min. Pellets were brought up in fresh RPMI-1640 and resuspended via sonication. Infectious EBs were then replated in a 96-well plate on fresh Cos-7 monolayers in a 2 fold dilution series. Re-infected monolayers were grown for 30 hpi prior to fixation with methanol before being stained with DAPI and Ctr MOMP Polyclonal Antibody, FITC (Fishersci). DAPI was used for visualization of host nuclei and for focusing our automated microscope. Anti-*Ctr* antibody was used for staining of re-infected inclusions for IFU counts. Inclusions were imaged using a Nikon Eclipse TE300 inverted microscope utilizing a scopeLED lamp at 470nm and 390nm, and BrightLine band pass emissions filters at 514/30nm and 434/17nm. An Andor Zyla sCMOS camera was used in conjunction with Micro-Manager software for image acquisition.

### Digital Droplet PCR

*Ctr* genomes were isolated using an Invitrogen PureLink genomic DNA minikit. Samples were then diluted as shown in (Sup Fig 4). DNA samples were added to ddPCR Supermix (Bio-RAD). Amplification was achieved using primer/probe sets for *nqrA* (origin) or *pyrG* (terminus). Droplets were generated using a Bio-RAD: QX200 AutoDG Droplet Digital PCR System. Data analysis was performed using Bio-RAD QX Manager 1.2 along with custom matplotlib, pandas, and seaborn python scripts.

### Fluorescent In-Situ Hybridization (FISH)

*Ctr* infected monolayers were fixed with 4% PFA for 10 minutes at room temp. Fixed coverslips were then permeabilized in 70% ethanol at -20 degrees Celsius overnight. Hairpin amplification was achieved using Molecular Instruments HCR FISH kit. The custom designed probes were ordered from Molecular Instruments and are listed in Table S2. Coverslips were mounted using MOWIOL and imaged using the CrestOptics X-Light confocal system.

### Click Chemistry

*Ctr* infections were treated with peptidoglycan incorporating reagent ethynyl-D-alanyl-D-alanine (EDA-DA) at a concentration of 1mM. Monolayers were fixed with 4% PFA for 10 minutes at room temp, fixed coverslips were permeabilized with 0.1% TritonX-100 prior to blocking with 1xPBS + 3% BSA. Click chemistry was carried out utilizing the Click-iT™ Plus Alexa Fluor™ 647 Picolyl Azide Toolkit according to the manufacturer’s instructions.

## Supplemental Material

**Figure S1:**
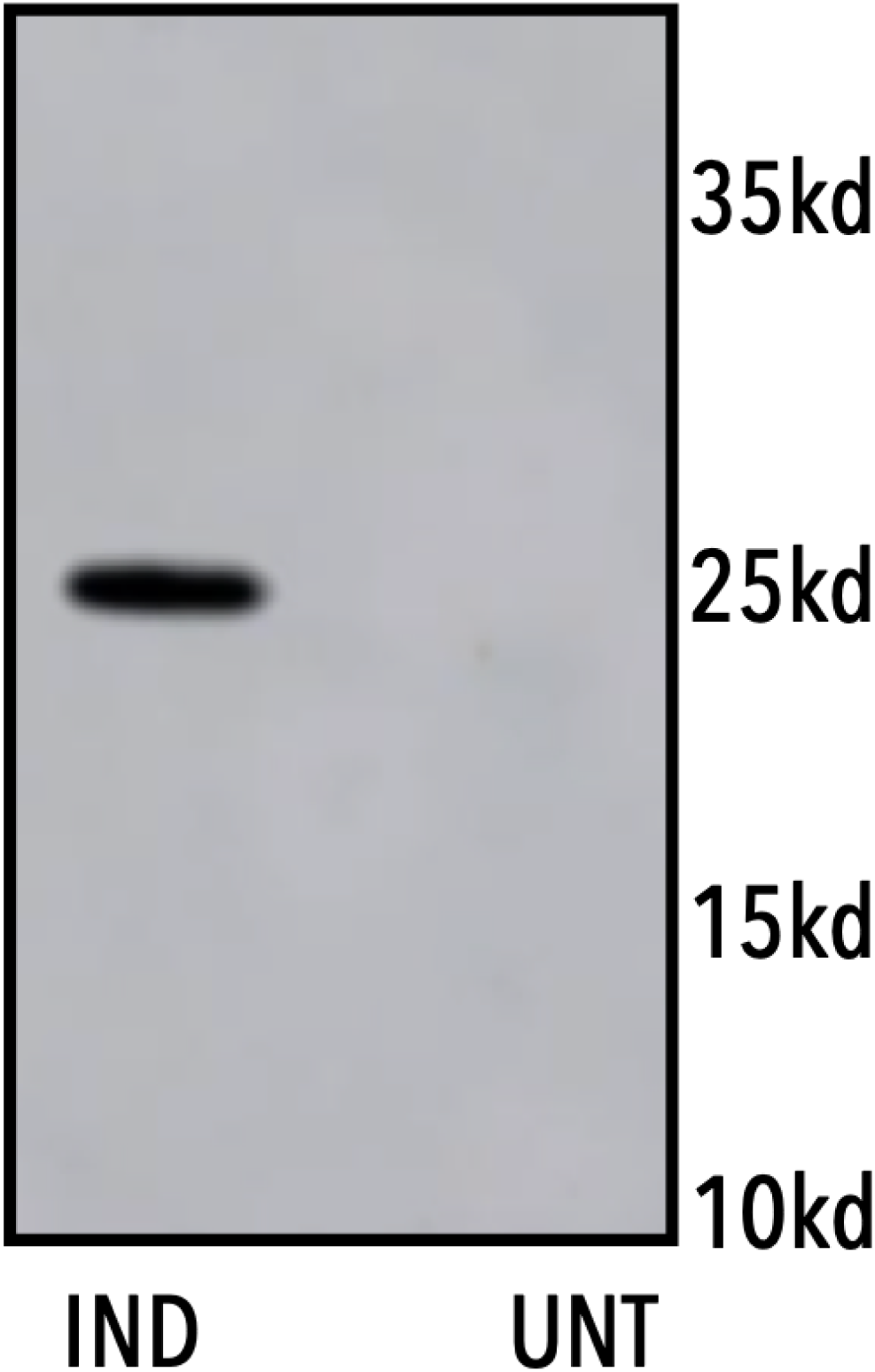
Western blot of Euo-FLAG. Anti-FLAG western blot of Cos-7 cells infected with L2-E-Euo-FLAG comparing Euo-FLAG expression in Tph treated and untreated cultures. Cells were induced or not with 0.5mM Tph at 16 hpi and proteins were harvested at 30 hpi, separated by PAGE, transferred to a nitrocellulose membrane and probed for the presence of the FLAG tag.

**Figure S2:**
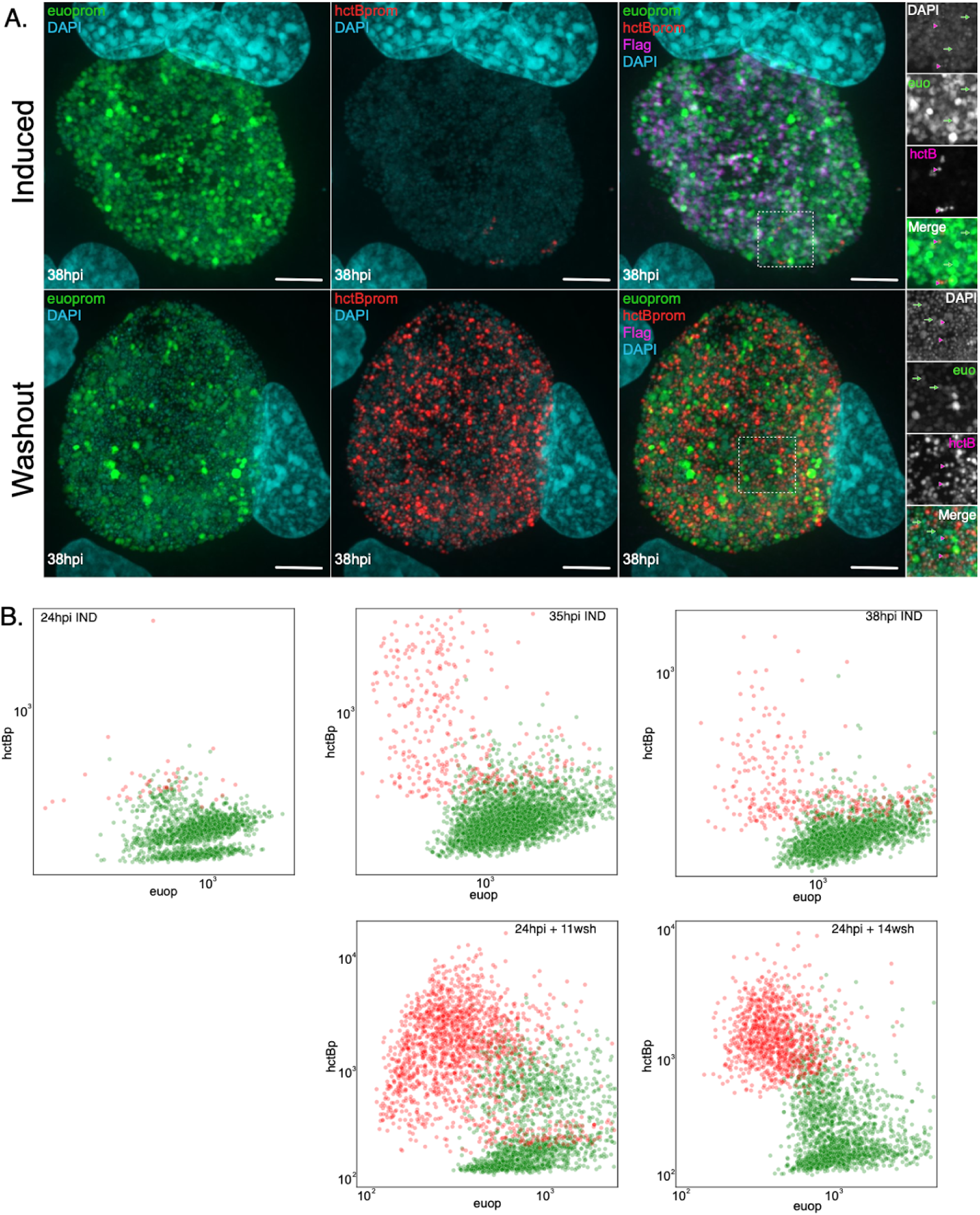
Reentry into the developmental cycle after inducer washout. A) Confocal micrographs of Cos-7 cells infected with L2-E-Euo-BmeC induced with 0.5mM TPH at infection. The 38 hpi inclusions in the induced control infected cells were imaged using confocal microscopy and had primarily *euo*prom+ cells (green) with few *hctB*prom+ cells (red). These cells were stained positive for Euo-FLAG (magenta). Tph was washed out at 24 hpi and the cells were fixed 14 hours later at 38 hpi, stained for Euo-FLAG expression and imaged using confocal microscopy. The washout inclusions had fewer *euo*prom+ cells (green) and an increase in *hctB*prom+ cells (red). These cells had essentially no staining for Euo-FLAG (megenta). Scale bar =15 µm. B) Cells were infected with L2-E-Euo-BmeC and induced with 0.5mM TPH at infection. Individual *euo*prom+ cells (green) and *hctB*prom+ cells (red) were identified using the Trackmate in FIJI and the fluorescence signal for both *euo*prom and *hctB*prom in individual chlamydial cells from five inclusions was determined. These values were plotted for each cell, green (*euo*prom+) and red (*hctB*prom+). At 24 hpi the inclusions contained primarily *euo*prom+ (green) fluorescent signal. Tph was removed, fluorescent intensity was determined for both *euo*prom+ (green) and *hctB*prom+ (red) cells at 11 and 14 hours post washout (hpw) and plotted. Red cells positive for *hctB*prom signal increased over time after washout.

**Figure S3.**
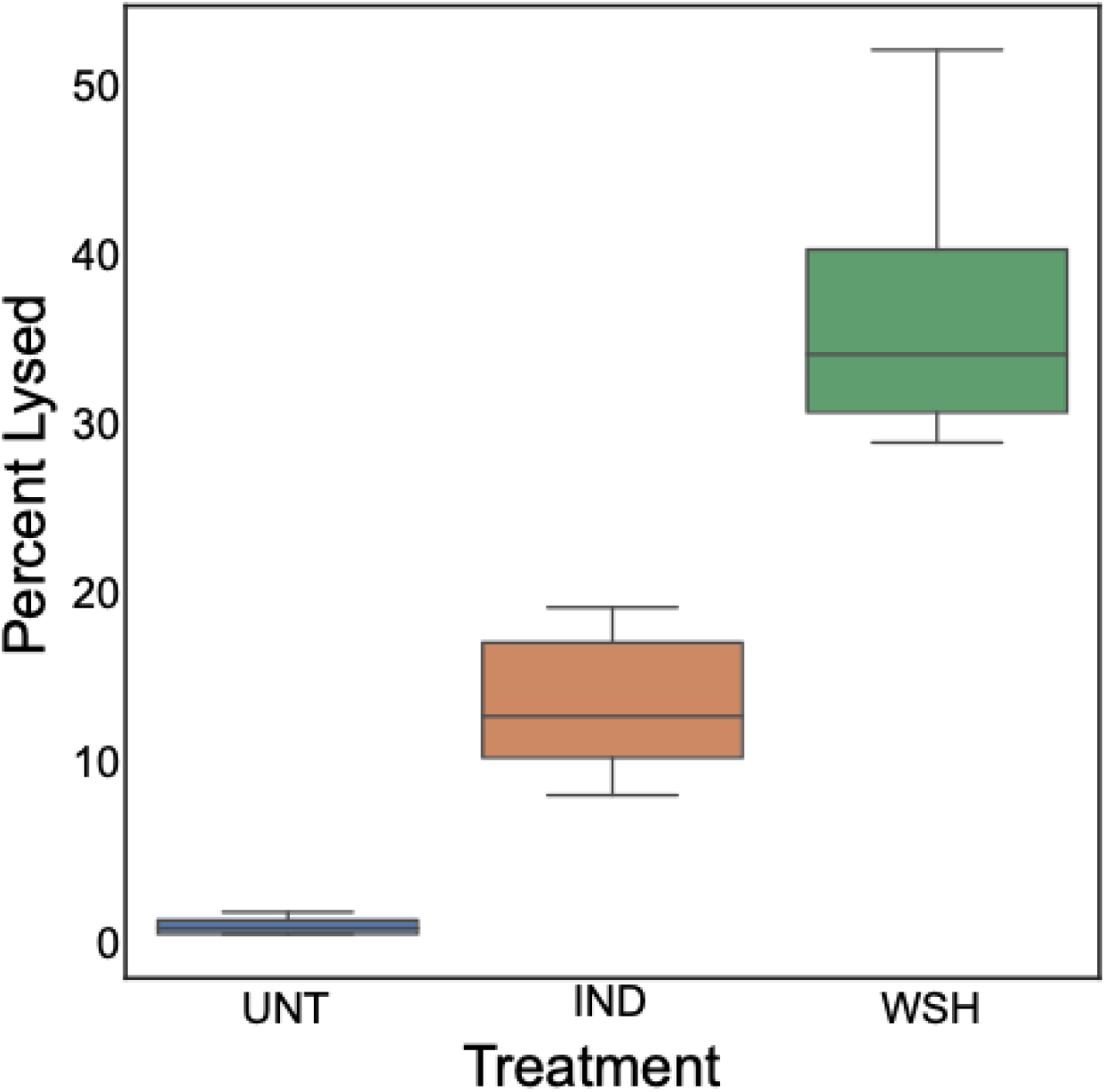
Quantification of inclusion/cell lysis after Tph washout. Cos-7 cells were infected with L2-E-Euo-BmEc and treated with Tph or vehicle only at infection. At 24 hpi Tph was washed out and the infected cultures were imaged every 30 minutes for expression of GFP (*euo*prom) and RFP (*hctB*prom). At 3 hours post washout the number of inclusions that visibly lysed was quantified for the untreated (UNT), Tph induced (IND) and Tph washout (WSH) cultures. Images were taken at 4x magnification. 8 FOV were counted per treatment with n > 100 inclusions counted per FOV.

**Figure S4:**
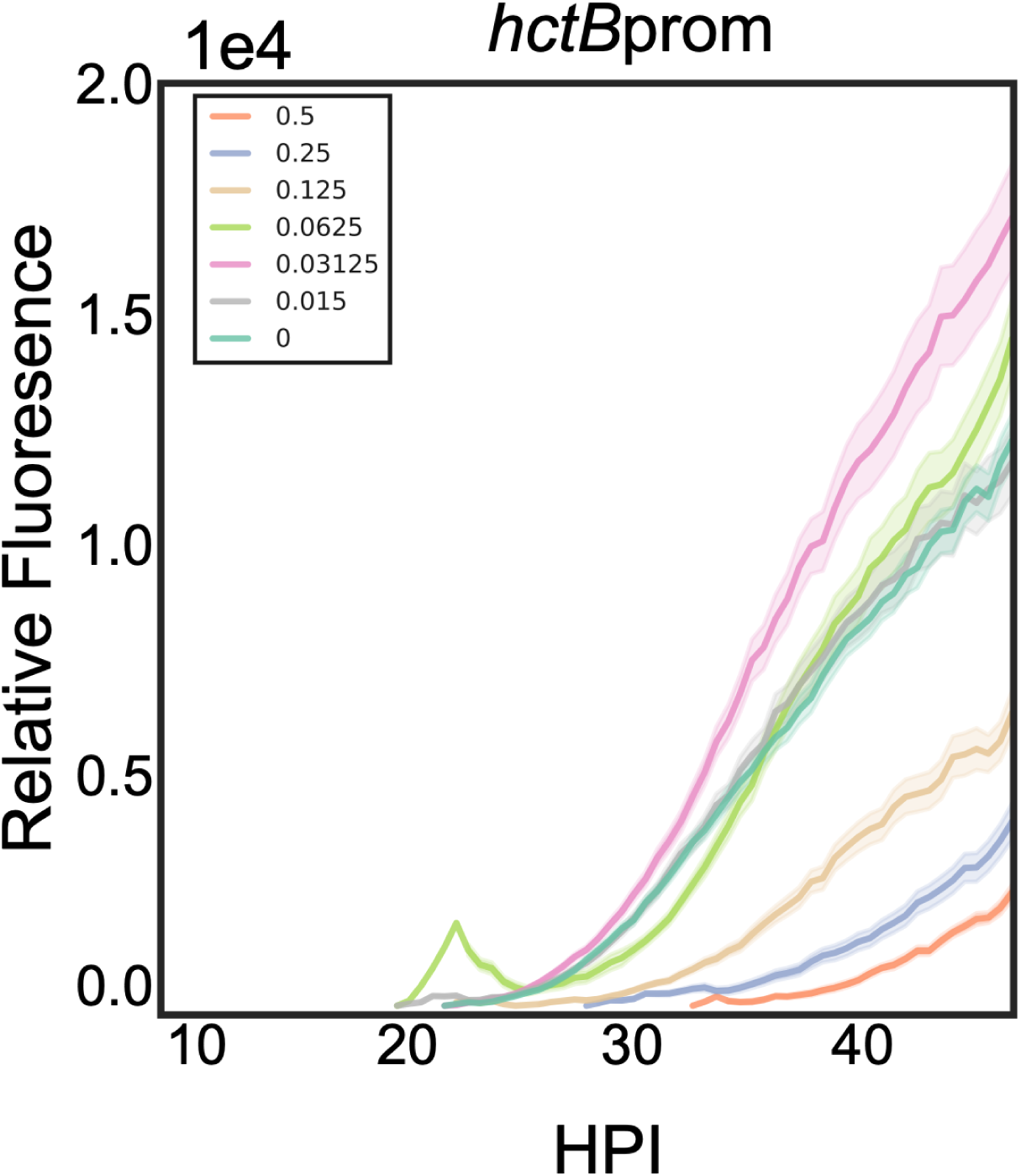
Kinetics of the developmental cycle of L2-euonprom-E-Euo-Bng induced for Euo expression with a range of Tph. Cos-7 cells infected with L2-*euo*nprom-E-Euo-Bng and were induced for Euo expression at 0 hpi with 0.5, 0.25, 0.125, 0.0625, 0.03125, 0.015 or Tph or vehicle only and assayed for *hctB*prom activity by live cell microscopy. Error cloud for fluorescent reporter represents SEM. n > 20 inclusions per treatment.

**Supplemental Table 1:**
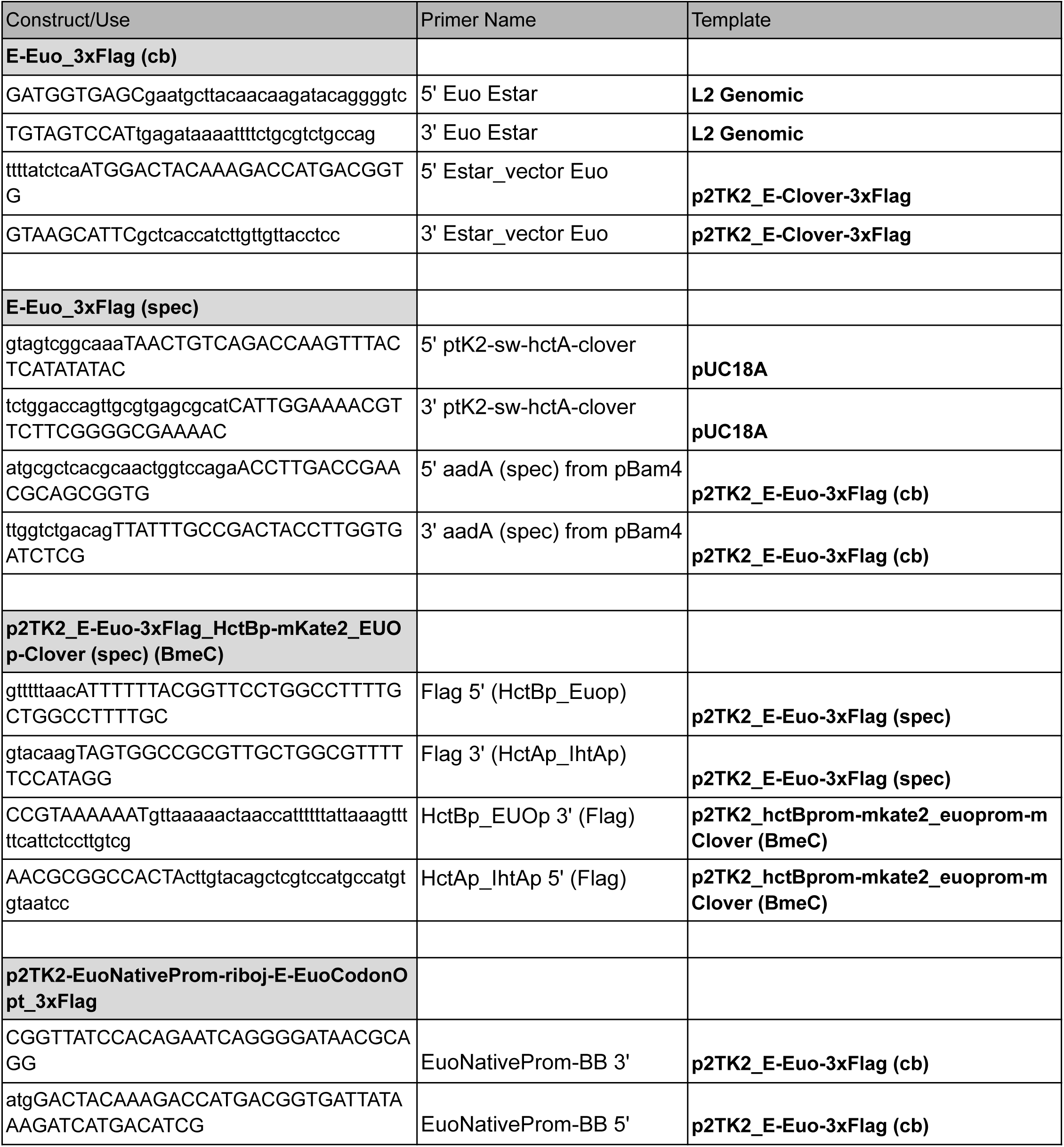
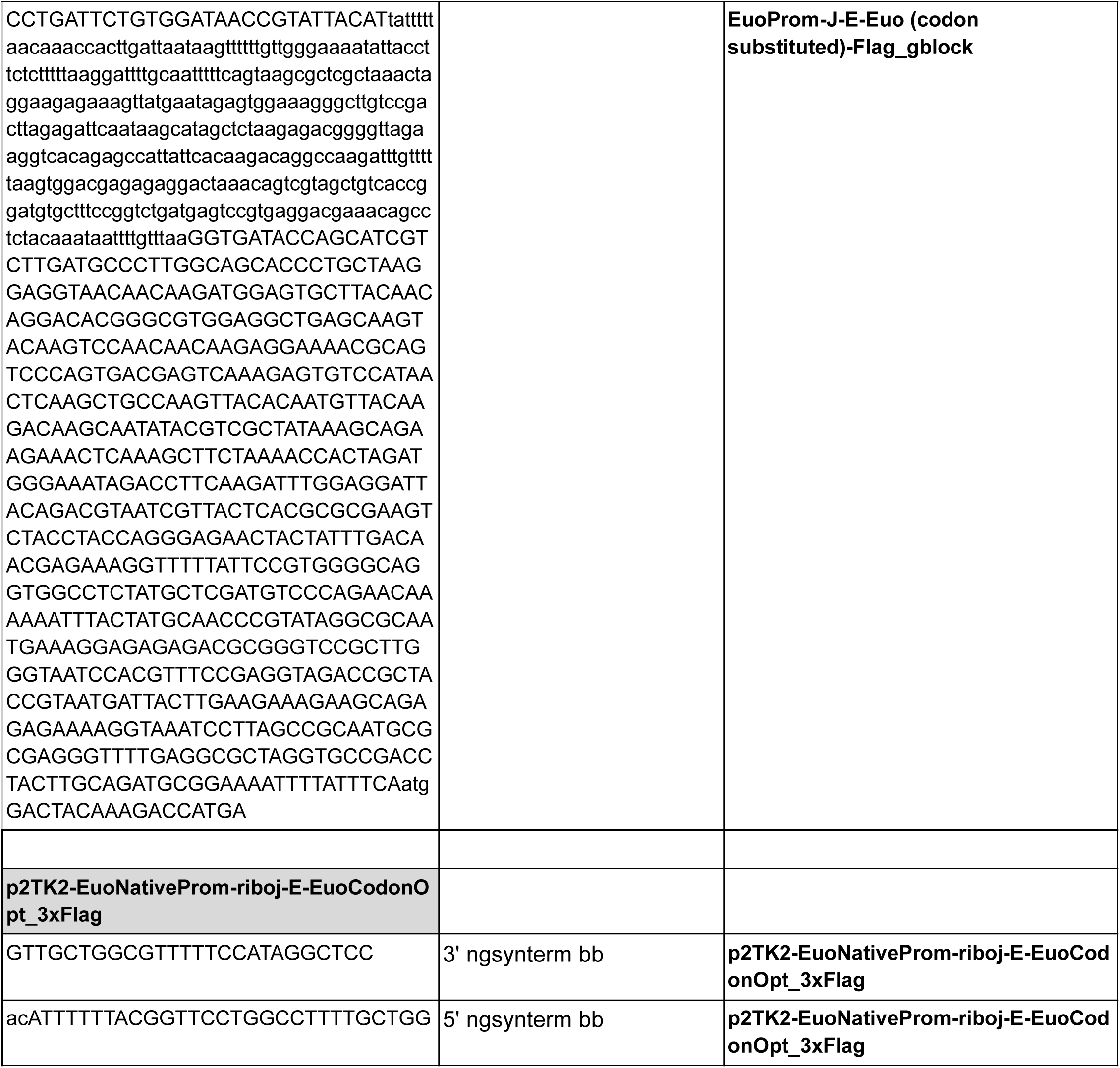
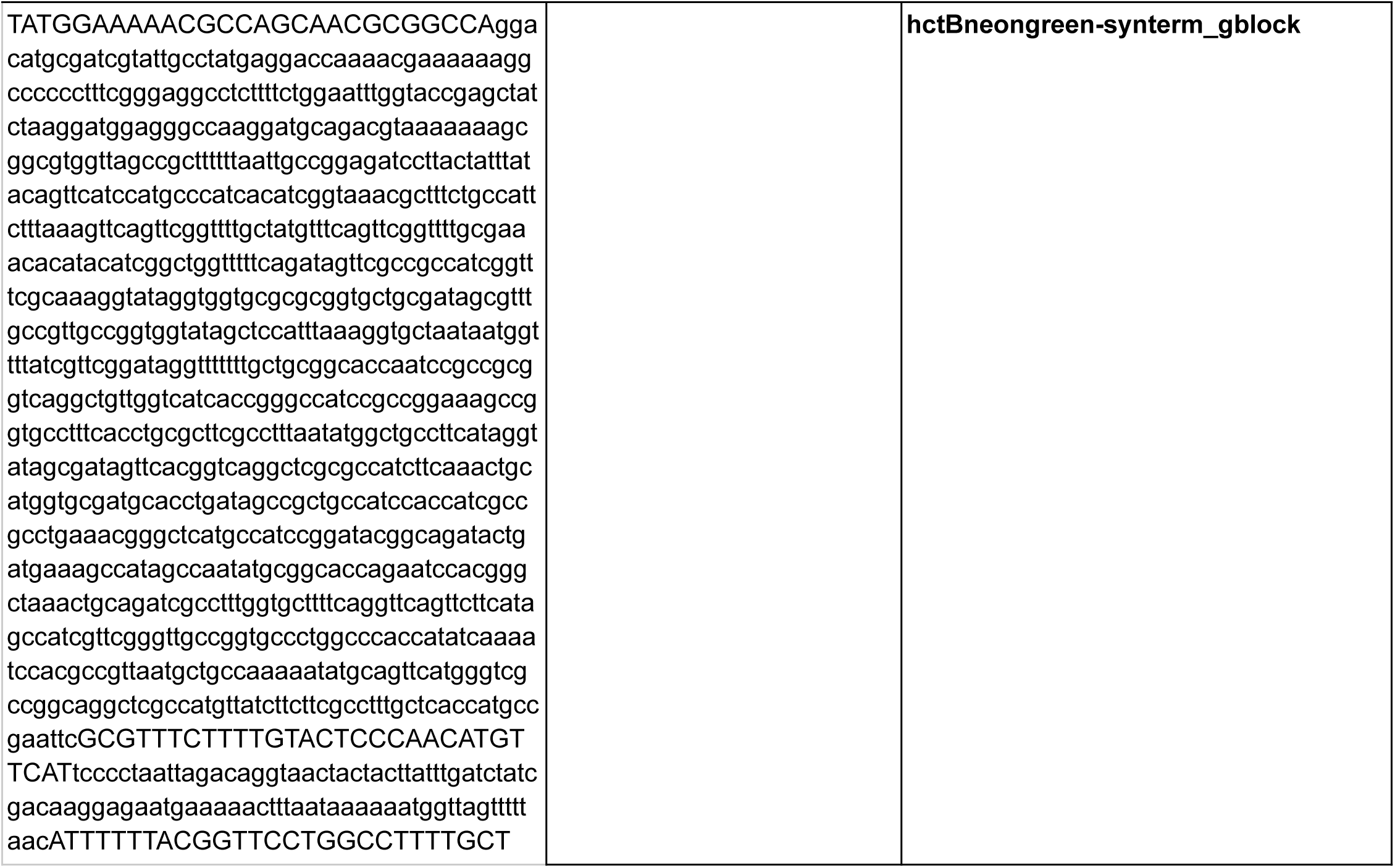
Plasmid and primer table

**Supplemental Table 2:** In-Situ Probes

| Gene | Seq Length | Seq Range |
| --- | --- | --- |
| euo | 739 | 203142 - 203880 |
| hctA | 561 | 895571 - 896131 |
| Tarp | 3072 | 849421 - 852492 |
| IncD-G | 1676 | 457415 - 459091 |

**Supplemental Movie 1**: Cos-7 cells infected with E-Euo-BmEc and induced for Euo-FLAG expression at infection with 0.5 mM Tph. At 24 hpi the infected cells were imaged every 30 minutes for expression of GFP (*euo*prom) and RFP (*hctB*prom) for an additional 56 hours.

**Supplemental Movie 2**: Cos-7 cells infected with E-Euo-BmEc and induced for Euo-FLAG expression at infection with 0.5 mM Tph. At 24 hpi Tph was removed and the infected cells were imaged every 30 minutes for expression of GFP (*euo*prom) and RFP (*hctB*prom) for an additional 56 hours.

